# Patient-informed CRISPR Screen Identifies FLNB as a Novel Congenital Heart Disease and Ciliopathy Gene

**DOI:** 10.1101/2025.10.14.682288

**Authors:** Angelo Arrigo, Venkatraman Rao, Aakrosh Ratan, Saurabh S Kulkarni

## Abstract

Heterotaxy (HTX) syndrome is a congenital disorder characterized by abnormal left-right organ placement, often leading to severe congenital heart defects (CHD). Despite advances in sequencing, many CHD/HTX-associated genes remain functionally unvalidated, hindering effective clinical diagnosis and management. Here, we leveraged a high-throughput CRISPR/Cas9 screening approach in the *Xenopus* model to rapidly evaluate candidate genes identified from whole-exome sequencing of human CHD patients. Our screen identified Filamin B (FLNB), an actin-binding protein previously linked to skeletal disorders but not to ciliopathies or CHD. We identified 5 probands with CHD/HTX, 3 with recessive, and 2 with damaging heterozygous mutations in FLNB. Disrupting FLNB in *Xenopus* reproduced key features of the human HTX phenotype, including defects in cardiac development and impaired motile cilia function. Rescue experiments confirmed the functional conservation of human FLNB, directly implicating actin cytoskeletal disruption in ciliogenesis and left-right patterning defects. Our results provide crucial evidence linking human FLNB dysfunction to ciliopathies and CHD/HTX.

## MAIN TEXT

Congenital heart disease (CHD) affects nearly 40,000 births per year in the United States^1^. Despite significant improvements in perinatal, operative, and perioperative care, the morbidity and mortality rates for infants with CHD remain unacceptably high^1^. Heterotaxy (HTX), also known as situs ambiguous, is one of the most severe forms of CHD, characterized by disrupted left-right (L-R) body asymmetry, often accompanied by severe organ abnormalities^2^. Despite advancements in genomic technologies, the genetic basis of CHD/HTX remains incompletely understood. Sequencing efforts in large cohorts have identified numerous candidate variants associated with CHD/HTX; however, validating the pathogenicity of these candidate variants remains a significant challenge^3-8^.

A recent large-scale genomic study identified multiple candidate genes involved in CHD and HTX^3^. However, systematic functional validation of these candidates in a model organism is important for differentiating pathogenic variants from false positives^9^. Current validation strategies, including mouse models, require significant time and resources, hindering rapid screening. Hence, there is a need for high-throughput, biologically relevant models to assess the functional consequences of these variants. To address this gap, we utilized the vertebrate model organism *Xenopus tropicalis*, which is well-established for studying developmental genetics due to its rapid embryogenesis and ease of genetic manipulation^10-14^. Given the prominent role of cilia dysfunction in both CHD and HTX^15,16^, *Xenopus* provides a uniquely powerful model for rapidly assessing ciliary function and functionally validating CHD/HTX candidate genes^10-12,17-20^.

Here, we implemented a targeted CRISPR/Cas9 loss-of-function (LOF) screening approach to rapidly assess the functional role of 18 genes identified as candidate HTX loci from the previously mentioned large-scale patient cohort^3^. Through systematic evaluation, we discovered that the disruption of Filamin B (*FLNB*), which encodes an actin-binding protein not previously associated with ciliopathies or CHD/HTX, recapitulated the HTX patient phenotypes in *Xenopus*. Our findings identify *FLNB* as a novel CHD/HTX and ciliopathy gene, demonstrating the efficacy and necessity of integrating patient-derived genomic data with high-throughput functional validation. By employing rapid CRISPR-based gene editing in *Xenopus*, our study provides immediate, impactful insights that directly link a candidate human genetic variant to a specific developmental phenotype.

A recent study performed whole-exome sequencing on 2,871 patients with congenital heart disease (CHD), identifying inherited recessive, dominant, and de novo variants^3^. Among these, 272 patients exhibited a bona fide heterotaxy (HTX) phenotype. To prioritize candidate genes contributing to HTX, we applied stringent variant-filtering criteria exclusively to HTX trio data, selecting for loss-of-function (LOF) recessive variants and damaging de novo variants. We excluded synonymous mutations, tolerated missense mutations, and LOF heterozygous variants, resulting in a final list of 57 prioritized genes. Of these genes, 11 had previously been established as associated with HTX; notably, 6 out of these 11 (∼50%) were cilia-related genes, highlighting the significant role of cilia in left-right (LR) patterning. This finding aligns well with previous mouse genetic screens for CHD, which identified approximately 50% of cilia-associated genes^15^.

To further explore candidate genes lacking prior HTX or ciliary associations, we analyzed the remaining 46 genes and identified *Xenopus* orthologs for 43. Based on ciliary associations predicted from the analysis of eight databases (Table S1), we selected 18 genes for a CRISPR-mediated loss-of-function (LOF) screen in *Xenopu*s embryos. In *Xenopus*, the orientation of the outflow tract (heart looping) is an established method for analyzing LR asymmetry^19,21^. The CRISPR screen identified several genes that are essential for establishing LR asymmetry (Figure 1A-C). We categorize genes that show more than 10% LR patterning defects, 6 genes (*FLNB, FRYL, GJA10, EEA1, MYO15A, ZC3H14*) in our screen, in *Xenopus* embryos as high-confidence. The remaining 11 genes are considered low-confidence genes. Filamin B (*flnb*) was the top hit, prompting further functional validation (Figure 1A).

**Figure 1.**
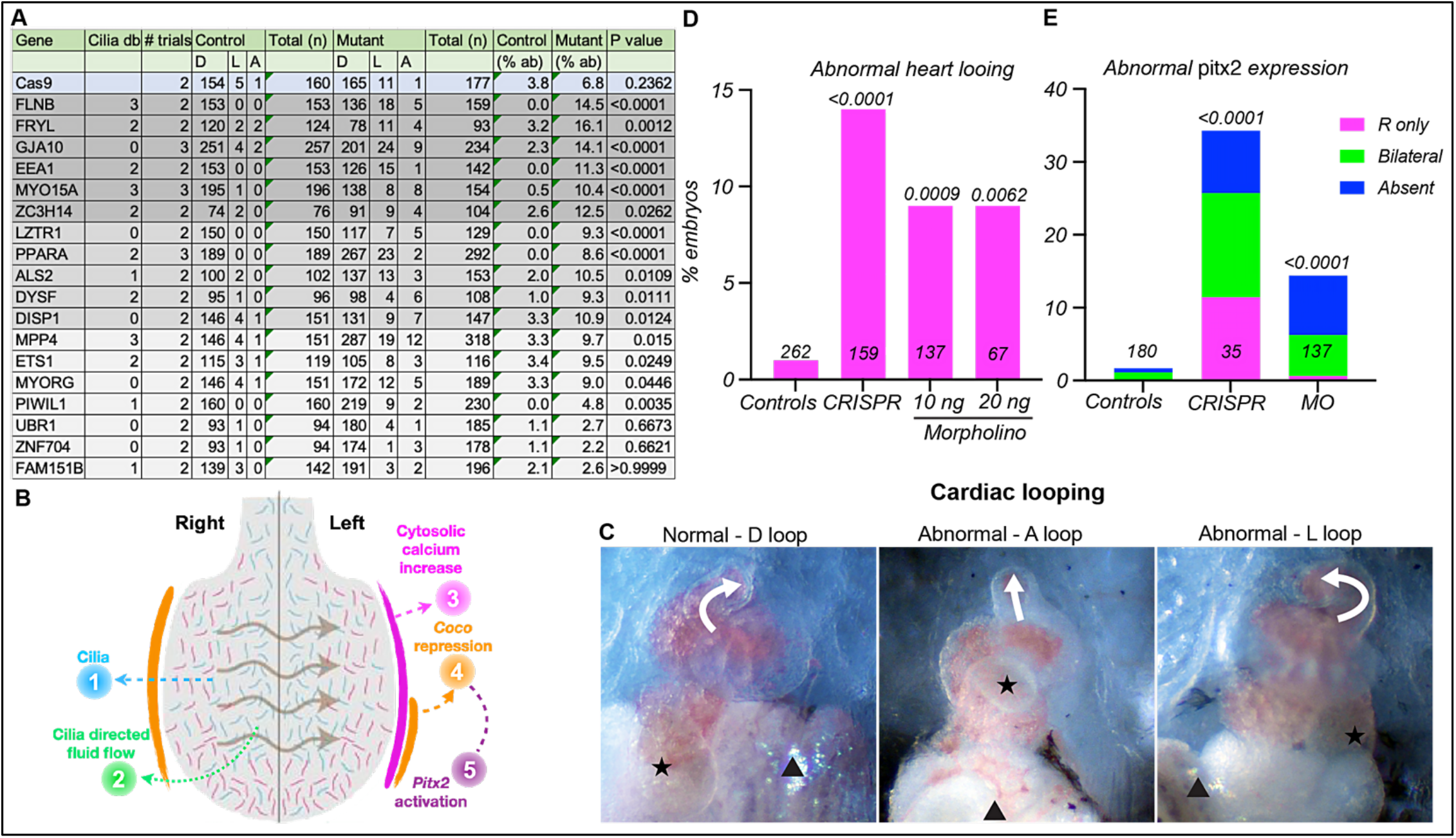
Flnb depletion results in LR patterning defects. (A) Eighteen predicted CHD/HTX-associated genes with varying previous associations with cilia were selected from whole-exome sequencing data of 2,871 patients for a CRISPR-mediated LOF screen in *Xenopus*. Flnb was the top candidate for the LR patterning defects identified in the screen. (B) Cilia in the LRO generate a leftward flow that sets off a gene cascade, repressing *coco*, and in turn activates *pitx2* in the left lateral plate, which alters organogenesis. (C) Normal heart looping is a result of left-sided *pitx2* expression. Abnormal heart looping in *Xenopus* embryos is indicative of an LR patterning defect leading to HTX and CHD. White arrows indicate the direction of the outflow track. Black star and arrowhead indicate the position of the gallbladder and gut in normal and abnormal conditions. (D, E) Control, CRISPR, 10 ng, or 20 ng morpholino-injected embryos were scored for (D) abnormal heart looping and (E) abnormal *pitx2* expression (20 ng MO only). Loss of Flnb causes both abnormal heart looping and abnormal *pitx2* expression. The number of embryos (n) scored for each experiment is indicated in the bar graph. The p-value is shown above each bar graph.

We validated CRISPR LOF results with morpholino oligo (MO)-based knockdown. Both 10 ng and 20 ng MO doses caused consistent heart looping defects, mirroring CRISPR results and human pathogenesis (Figure 1C, D). These defects resemble human HTX, supporting Flnb as a vertebrate HTX gene.

To understand the molecular underpinnings of the heart looping defects, we performed *in situ* hybridization for *Pitx2*, a marker of LR asymmetry^21^. *Pitx2* is normally left-sided; controls showed normal expression, while *flnb* mutants and morphants exhibited misexpression (Figure 1E). Since motile and sensory cilia within the left–right organizer (LRO) drive the molecular cascade that restricts *pitx2* to the left, we next examined LRO morphology and cilia formation. Loss of FLNB resulted in a markedly abnormal LRO: morphants exhibited a narrower structure with a higher length-to-width ratio than controls (Figure 2A, B). This was accompanied by a significant reduction in cilia number (Figure 2C). Analysis of cilia length distributions revealed that controls displayed a broad spectrum, ranging from short (<2.5 μm) and intermediate cilia (∼2.5-4.5 μm) to long cilia reaching 7 μm. By contrast, morphants showed a sharp peak at 2.5–3.5 μm followed by a steep decline, consistent with impaired cilia elongation (Figure 2D, F). This loss of broad distribution is also reflected by a significantly smaller interquartile range in morphants compared to controls (Figure 2E). Together, these findings indicate that Flnb is essential for proper LRO architecture and cilia development, with its loss leading to depletion of short, likely sensory cilia at the LRO margins and long motile cilia at the center.

**Figure 2.**
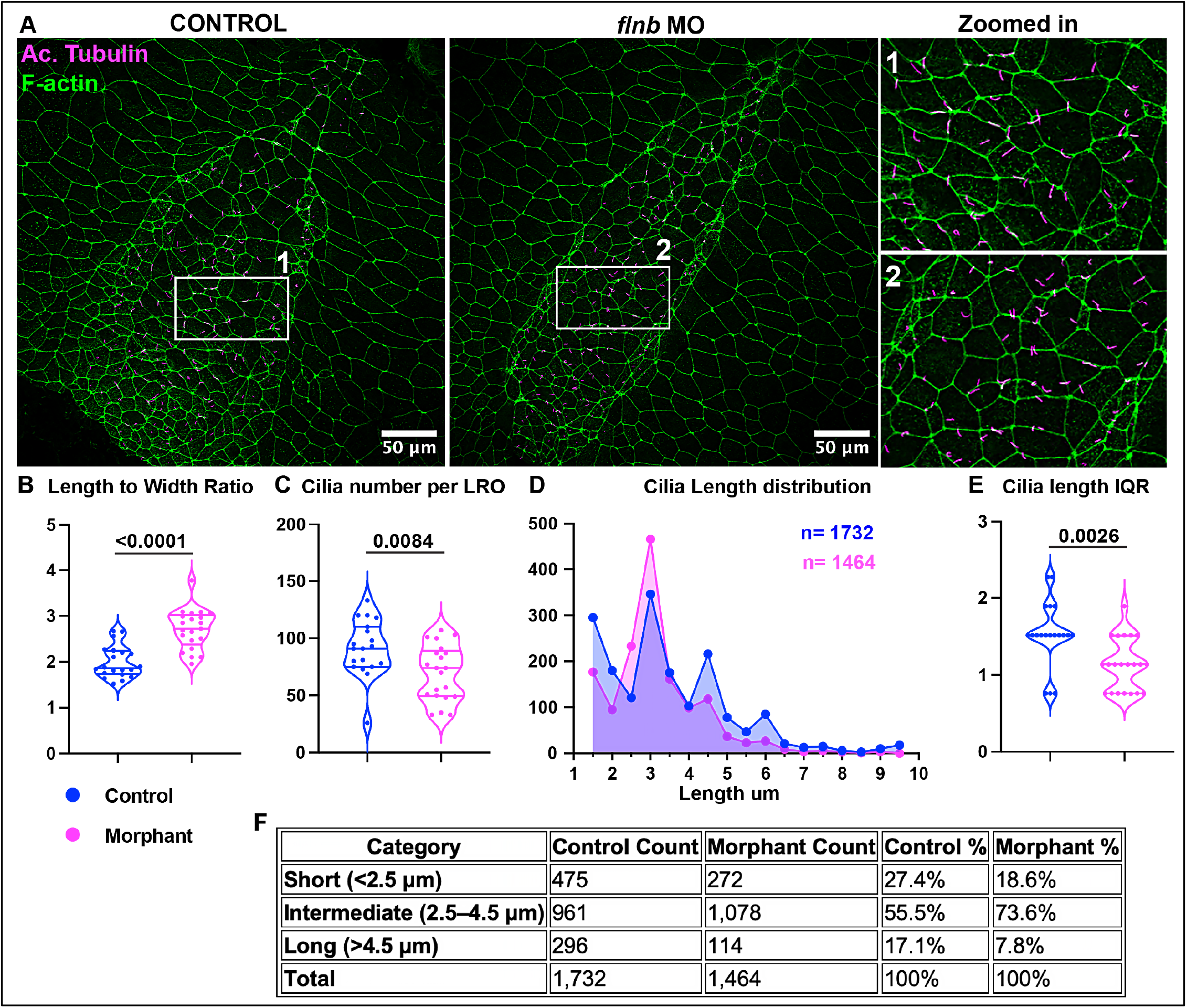
Flnb depletion leads to defects in the left-right organizer and in ciliogenesis. (A) Left-right organizer (LRO) in control and morphant *Xenopus* embryos at embryonic stage 16. White boxes indicate the zoomed-in region of the LRO. (B) Statistical comparison of the Length-to-width ratio of LROs in control and morphants. (C) Statistical comparison of cilia number between control and morphants. (D) Histogram showing cilia length distribution in control and morphants. (E) Inter-quartile range (25%-75%) showing distribution of cilia length in controls and morphants. (F) Table showing. Number of cilia in each category of cilia length: short, medium, and long. n = 19 and 21 LROs for controls and morphants, respectively, from 2 trials. The p-value is shown above each bar graph

To determine whether FLNB variants were identified in other CHD phenotypes, we examined the remaining proband dataset from the study. ^3,22^. We identified four additional probands: two with damaging recessive mutations (conotruncal defects and left ventricular outflow tract obstruction) and two with loss-of-function heterozygous mutations (conotruncal defects and other cardiac defects) (Table 1), further supporting the evidence that FLNB is crucial for cardiac development and LR patterning and extend the previous findings that link FLNB to Larsen syndrome.

**Table 1.** Summary of additional FLNB variants identified in non-heterotaxy CHD probands from the Jin et al. (2017) dataset, highlighting damaging recessive and heterozygous loss-of-function mutations associated with diverse cardiac phenotypes. The new study, by Dong et al. (2025), listed the same patients and identified FLNB as a candidate CHD gene after additional analysis.

| List of damaging *FLNB* recessive genotypes in CHD case cohort |  |  |  |  |
| --- | --- | --- | --- | --- |
| Blinded ID | Cardiac category | Type | AA chnage | Phenotype |
| 1-04709 | CTD | CmpHet | K137R/D623V | Right ventricular volume overload \| tetralogy of fallot |
| 1-02990 | LVO | CmpHet | P157L/F1411L | Aortic stenosis - valvar \| bicommissural (bicuspid, bileaflet, bifoliate) aortic valve \| left aortic arch with normal branching pattern |
| 1-07394 | HTX | CmpHet | T1814M/V2096M | Double outlet right ventricle \| heterotaxy \| hypoplastic mitral valve \| partially anomalous pulmonary veins \| sda \| single ventricle in heterotaxy syndrome \| single ventricle, right, double inlet right ventricle \| univentricular, double inlet atrioventricular connection |
| Loss-of-function *FLNB* heterozygous mutations in cases |  |  |  |  |
| 1-00366 | CTD | frameshift_deletion | p.C450fs | Congenital pulmonary valve abnormality \| dilated right pulmonary artery \| hypoplastic pulmonary annulus \| left aortic arch \| main pulmonary artery abnormality \| tetralogy of fallot \| ventricular septal defect, malalignment |
| 1-02266 | OTHER | frameshift_deletion | p.I370fs | Tricuspid atresia + no pulmonary stenosis and large vsd (ic) \| ventricular septal defect, multiple \| ventricular septal defect, muscular |

We used a deep learning model, AlphaMissense, to further assess the predicted impact of recessive missense variants in probands affecting FLNB function (Table 1, Figure S1-S3)^23^. For each of the three patients, we identified at least one missense mutation as non-benign (Pathogenicity score > 0.34). Analysis of all possible missense variants in the gene showed that more than 80% of possible missense mutations overlapping the actin-binding domain and the disordered region were predicted to be pathogenic. We also analyzed the likely pathogenic and pathogenic FLNB missense variants from ClinVar that have been implicated in Larsen syndrome and other FLNB-related disorders using AlphaMissense. We found that the majority of the reported ClinVar variants are located within the actin-binding domain and the repeat region between the Hinge and the FBLP1-interacting domain (Figure S1).

Motile cilia play a pivotal role in establishing left-right (LR) asymmetry and mucociliary clearance.^11,16,24-26^. Therefore, patients with CHD and HTX show a high prevalence of respiratory complications^27-31^. Moreover, a recent study of the single-cell RNA sequencing of human lungs showed high FLNB expression in multiciliated cells^32^. Therefore, we investigated FLNB’s function in motile cilia assembly and function using the *Xenopus* embryonic epidermal multiciliated cell (MCC) model^11,33^. To this end, we examined cilia-driven extracellular fluid flow over the surface of the *Xenopus* embryo to assess cilia function in MCCs^17,33^. We visualized the flow by adding latex microspheres (beads) to the culture media, followed by time-lapse imaging of bead movement. In wild-type (WT) embryos, cilia-generated flow was brisk (Figure 3B). In contrast, *flnb* morphants displayed significantly impaired or absent fluid flow, suggesting a dramatic loss of cilia (Figure 3B). Detailed immunofluorescence analysis of *flnb* morphants showed that the MCCs were notably smaller, with a dramatic loss of cilia, specifically fewer cilia as measured by reduced acetylated tubulin staining (Figure 3A, C, D). MCCs normally organize a dense F-actin network on their apical surface, which is crucial for cell shape and basal body organization^17^. *Flnb* morphants showed significant disruption of apical F-actin organization, as measured by reduced cortical actin and medial F-actin enrichment in MCCs (Figure 3C, D). Consistently, *flnb* morphants also showed defects in apical expansion and cell shape. (Figure 3C, D). Taken together, these results bolster the evidence that FLNB may play a crucial role not only in establishing LR patterning but also in mucociliary clearance via its role in cilia. These findings were interesting because some cases of Larsen syndrome also show hydrocephalus, which is also a ciliopathy^25,34-36^.

**Figure 3.**
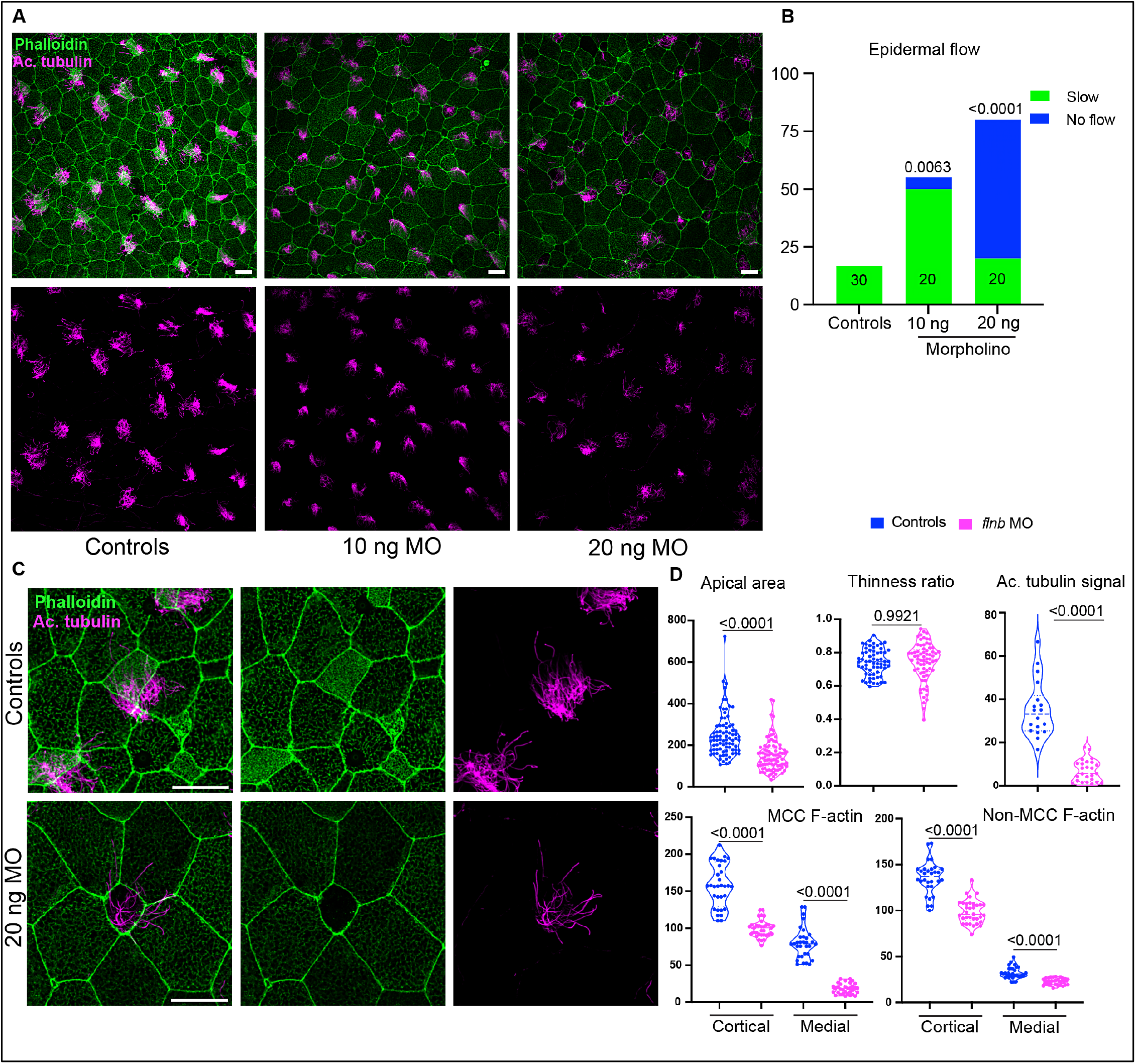
Flnb depletion results in dysfunctional multiciliated cells with aberrant gross morphology. (A) Control and *flnb* morpholino-injected embryos were stained with anti-acetylated α-tubulin antibody (magenta) and phalloidin (green) to label cilia and F-actin, respectively. (B) Mucociliary clearance in control and Flnb-depleted embryos was evaluated by measuring the rate of flow of beads over the surface of the embryos. The flow rate was categorized as either normal, slow, or none. *Flnb* morphants showed significantly slow or no flow. (C) Higher magnification of MCC morphology showing loss of apical F-actin, reduced cilia, and decreased α-tubulin intensity in flnb morphants. (D) Quantification of MCC morphology parameters: apical area, thinness ratio, acetylated α-tubulin signal, and cortical and medial F-actin intensity. Cortical and medial F-actin intensity was also assessed in non-MCCs. n = 30-80 cells from 9 embryos across three trials.

Next, we investigated the subcellular localization of human FLNB protein in *Xenopu*s embryos. Injection of GFP-tagged human FLNB (hFLNB-GFP) mRNA into the embryos revealed colocalization of FLNB-GFP with apical and sub-apical F-actin networks in MCCs, directly linking FLNB to the cytoskeletal integrity required for ciliogenesis (Figure 4A, B) ^17,37,38^. Finally, we demonstrated functional conservation by rescuing the *flnb* morphant phenotype with human FLNB (Figure 4C). Specifically, injecting hFLNB-GFP mRNA into one cell of a four-cell stage FLNB morphant embryo successfully restored both medial F-actin architecture and cilia formation in targeted MCCs, not only confirming the specificity of the MO depletion but also the critical role of human FLNB in F-actin organization and motile ciliogenesis (Figure 4C-E).

**Figure 4.**
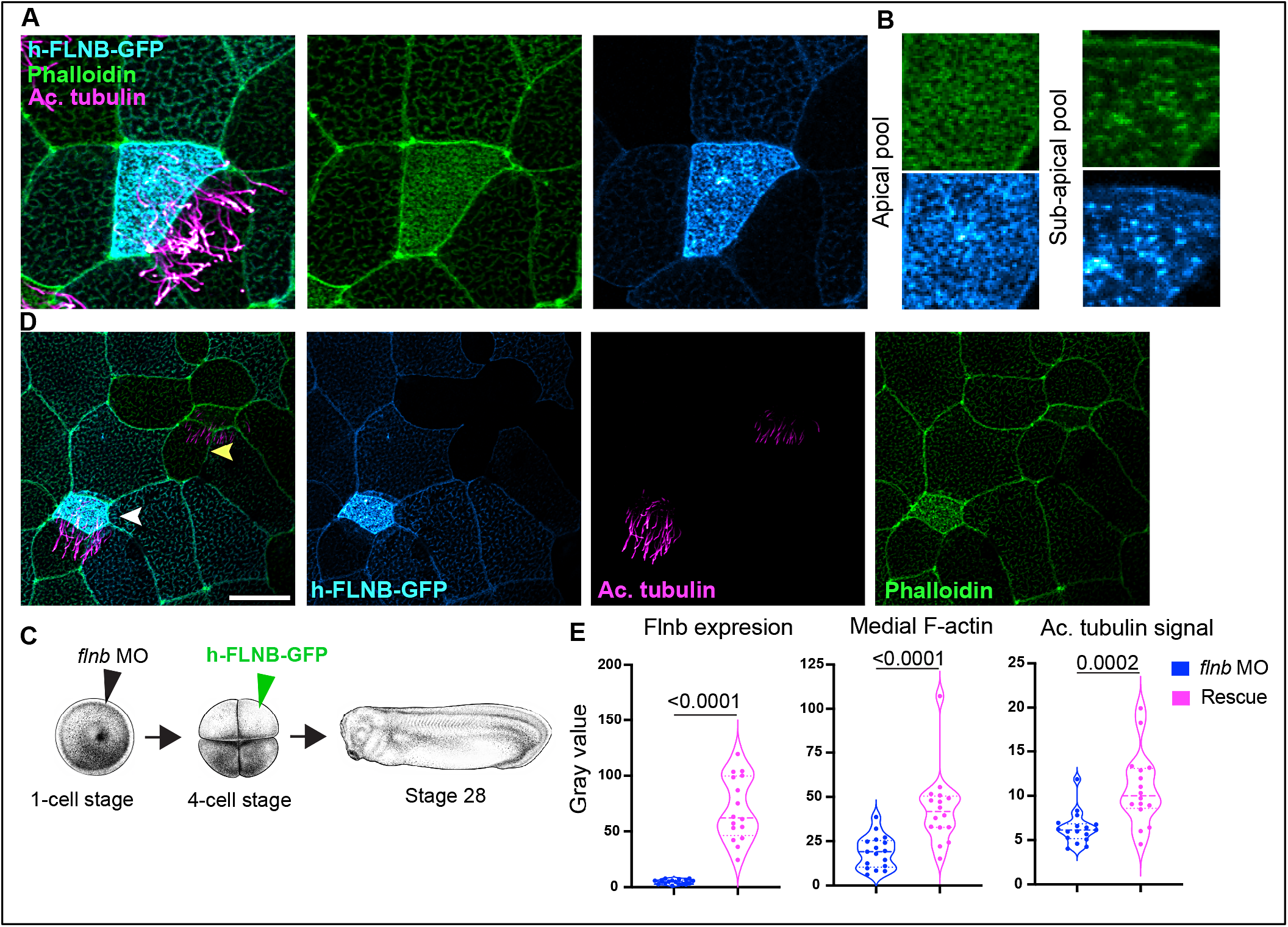
h-FLNB localizes to both the apical and subapical pool of F-actin and rescues the *flnb* morphant phenotype of MCCs. (A-B) h-FLNB-GFP was injected into WT fertilized eggs, and embryos were grown to stage 28, fixed, and stained to analyze the localization of h-FLNB in MCCs. h-FLNB-GFP (cyan) localizes to the apical meshwork and subapical pool of F-actin in MCCs. (C-E) To assess the functionality of the h-FLNB-GFP construct in *Xenopus, flnb* morphants were injected (rescued) at the 4-cell stage with h-FLNB-GFP, grown to stage 28, fixed, and stained. Rescued cells were marked by GFP expression (cyan). Flnb expression, medial F-actin, and acetylated tubulin intensity were quantified in morphant and rescue cells. H-FLNB-GFP rescued medial F-actin and cilia intensity. n=16-17 MCCs from 7-9 embryos across three trials.

A central mechanistic insight from our study is that FLNB links cytoskeletal integrity to motile ciliogenesis. Our data show that loss of Flnb disrupts apical F-actin organization in multiciliated cells, resulting in impaired apical expansion, reduced cilia number, and defective fluid flow. These findings place FLNB among a growing group of cytoskeletal regulators that govern ciliogenesis^12,17,38-45^, supporting the concept that ciliary dysfunction in CHD and HTX is not restricted to mutations in canonical ciliopathy genes, but can also arise from perturbations in actin-ciliary crosstalk. Consistent with this model, analysis of FLNB variants revealed pathogenic enrichment in domains mediating actin binding and cytoskeletal remodeling, directly aligning human genetic evidence with our mechanistic observations in *Xenopus*. Together, these results suggest that FLNB dysfunction compromises left– right patterning by impairing the actin–ciliary interface that is essential for coordinated motile cilia function.

Although FLNB knockout (KO) mice have not been reported to display laterality defects, Zhou et al. observed greater than 97% embryonic lethality, underscoring the critical requirement for FLNB in early development^46^. More recently, Huang et al. described non-syndromic orofacial clefts in their KO models^47^, phenotypes not noted by Zhou et al.^46^, raising the possibility that certain features were not examined in the earlier study or that phenotypic variability exists even within murine systems. Notably, neither study systematically assessed cardiac development, highlighting that the absence of reported findings should not be conflated with the absence of phenotype. Such discrepancies between mice and humans may also reflect species-specific regulatory programs, compensatory mechanisms, or strain-dependent genetic modifiers that can mask disease-relevant manifestations in mice. In particular, functional redundancy among filamin family members raises the possibility that FLNA partially compensates for the loss of FLNB in mice, thereby masking phenotypes that are more readily uncovered in *Xenopus* or human contexts, especially in cardiac development, where FLNA is critical and its loss is associated with cardiac malformations and heterotaxy-related defects^48-51^. Therefore, while the absence of phenotype in the mouse model should be noted, it does not undermine the strength of our *Xenopus* and human clinical data; instead, it highlights the importance of studying gene function across multiple model organisms to capture human pathogenesis.

Our findings also have important clinical implications. Patients with CHD and HTX frequently experience respiratory complications, including recurrent infections and impaired mucociliary clearance^27-31,52,53^. By demonstrating that FLNB is required for motile ciliogenesis and function in *Xenopus* and given its strong expression in human lung multiciliated cells, our results provide a plausible mechanistic explanation for these comorbidities. Moreover, the identification of FLNB mutations across diverse CHD subtypes, together with its established role in Larsen syndrome, suggests that FLNB-related disorders extend beyond skeletal disease to encompass a broader ciliopathy spectrum. In conclusion, our study identifies FLNB as a novel CHD and HTX gene, integrating patient genomic data with high-throughput *Xenopus* functional validation to reveal its essential role in cytoskeletal organization, ciliary function, and cardiac development. Importantly, our work demonstrates that *Xenopus* is an effective and relevant model for functional genomics, enabling rapid prioritization of candidate genes from large-scale patient sequencing efforts. By bridging the gap between human genomics and developmental biology, this study sets the stage for better diagnostic classification, deeper understanding of mechanisms, and ultimately, targeted therapies for patients with CHD and heterotaxy.

## DATA AVAILABILITY

This study does not include any data deposited in external repositories.

## ACKNOWLEDGMENTS

We thank Dr. Karen Hirsch for providing access to the confocal microscope. We are grateful for the NIH grant, NIGMS R35GM146856, and Saving Tiney Hearts Society grant awarded to Saurabh Kulkarni and NICHD R03HD112688, awarded to Saurabh Kulkarni and Aakrosh Ratan.

## AUTHOR CONTRIBUTIONS

AA: Investigation, Visualization, and Manuscript original draft writing and revisions.

VR: Investigation, Analysis, and Manuscript original draft writing and revisions.

AR: Bioinformatic analysis of the FLNB and pathogenicity predictions, and Manuscript revisions.

SSK: Conceptualization, Methodology development, Investigation, Visualization, Supervision, and Manuscript writing and revisions.

