## Supplemental Information for "Patient-informed CRISPR Screen Identifies FLNB as a Novel Congenital Heart Disease and Ciliopathy Gene"

### MATERIALS AND METHODS

#### Animal husbandry and in vitro fertilization

Sex as a biological variable is not relevant, as frog embryos are analyzed within 24-72 hours after fertilization. Frogs (*Xenopus tropicalis*) were bred and housed in a vivarium using protocols (ACUC# 4295) approved by the University of Virginia Institutional Animal Care and Use Committee (IACUC). Embryos needed for experiments were generated using in vitro fertilization, as described previously [54,55]. Briefly, the testes from male frogs were crushed in 1× MBS (pH - 7.4) with 0.2% BSA and added to eggs obtained from the female frogs. After 3 min of incubation, freshly made 0.1× MBS (pH - 7.8) was added, and the eggs were incubated for 10 more minutes till contraction of the animal pole of the eggs was visible. The jelly coat was removed using 3% cysteine in 1/9× MR solution (pH 7.8–8.0) for 6 min. The embryos were microinjected with morpholino/RNA and raised at 25 °C or 28 °C till they reached appropriate stages for heart looping, fixation and staining, or bead flow analysis. Embryos were staged as described previously[56]. The fertilized embryos for microinjection were chosen randomly.

#### Pathogenicity predictions

We used AlphaMissense to predict the pathogenicity scores for all possible missense variants in the human FLNB gene (NM_001457.4). Domain information for the gene was obtained from UniProt[57], and overlapping pathogenic and likely pathogenic mutations were retrieved from ClinVar[58].

#### RNA, morpholino, and microinjections

The human FLNB (NM_001457.4) plasmid was purchased from GeneCopoeia and subcloned into a pCS2+ destination vector using Gateway cloning. The RNA used in this study was generated by linearizing the plasmid and in vitro transcribed using the mMessage and mMachine SP6 transcription kit, followed by purification with the Zymo RNA purification kit. hFLNB-GFP RNA was microinjected at the 1-cell stage or 4-cell stage using glass needles mounted on the Pico-liter microinjection system (Warner Instruments). The FLNB translation-blocking morpholino was designed by GeneTools and injected (concentrations: 10 ng and 20 ng) at the 1-cell stage. FLNB sgRNA GGGAGGTGCTCAGTCAGAAA was designed using CRISPRscan to target the first exon and injected at the 1-cell stage.

**In situ hybridization**

*X. tropicalis* were collected at stage 28 and fixed in MEMFA (100 mM MOPS, 2 mM EGTA, 1 mM $MgSO_4$, 3.7% formaldehyde) for 1 hour. In situ hybridization was performed according to standard protocol[59] using an antisense probe for *pitx2*.

**Immunofluorescence staining and imaging**

The *X. tropicalis* embryos used for the study were fixed at stage 28 with 4% paraformaldehyde (PFA). After fixation, the embryos were washed three times with PBST (1× PBS with 0.2% Triton X-100) for 10 minutes each and then incubated in a blocking solution (3% BSA in PBST) for 1 hour. Appropriate antibodies were added to the embryos, incubated for 1 hour at room temperature, and then washed three times for 10 minutes each with PBST. A conjugated secondary antibody was used to stain embryos for 1 h. The embryos were washed three times with PBST (PBS + Triton), stained with phalloidin in PBST for 45 minutes, and then washed once with PBS after staining. The embryos were mounted and imaged. Confocal imaging was performed using the Leica DMi8 SP8 microscope with a 40x oil immersion objective (1.3 NA). Images were captured at 1× or 4× zoom, adjusted for brightness and contrast, analyzed in Fiji, and assembled in Adobe Illustrator software.

**Bead flow analysis**

Embryos were raised to stage 28 and anesthetized with benzocaine (0.05% in 1/9x MR). One microliter of 5.19 μM red beads (Bangs Laboratories, DSCR006) was placed at the anterior end of the embryo and visualized under a dissecting scope. Both control and morphant embryos were visually and qualitatively scored (fast, slow, none) for the speed of bead flow from the anterior to the posterior of the embryo.

**Protein variant modeling**

The FLNB protein sequence was retrieved from UniProt (O75369) and used for AlphaFold 3 modeling. For comparison, experimentally resolved structures of different FLNB domains were obtained from the RCSB PDB (Accession numbers: 2WA5, 2DIC, 2EE9, 2DI8). The corresponding amino acids from the wild type were replaced with those from the patient mutation for modeling. Since no structure for Filamin B repeat domain 4 (wild type) was available, that region of the protein (amino acids 544-636) was modeled using AlphaFold3[60]. All structures were visualized using UCSF ChimeraX[61-63].

### SUPPLEMENTAL FIGURE LEGENDS

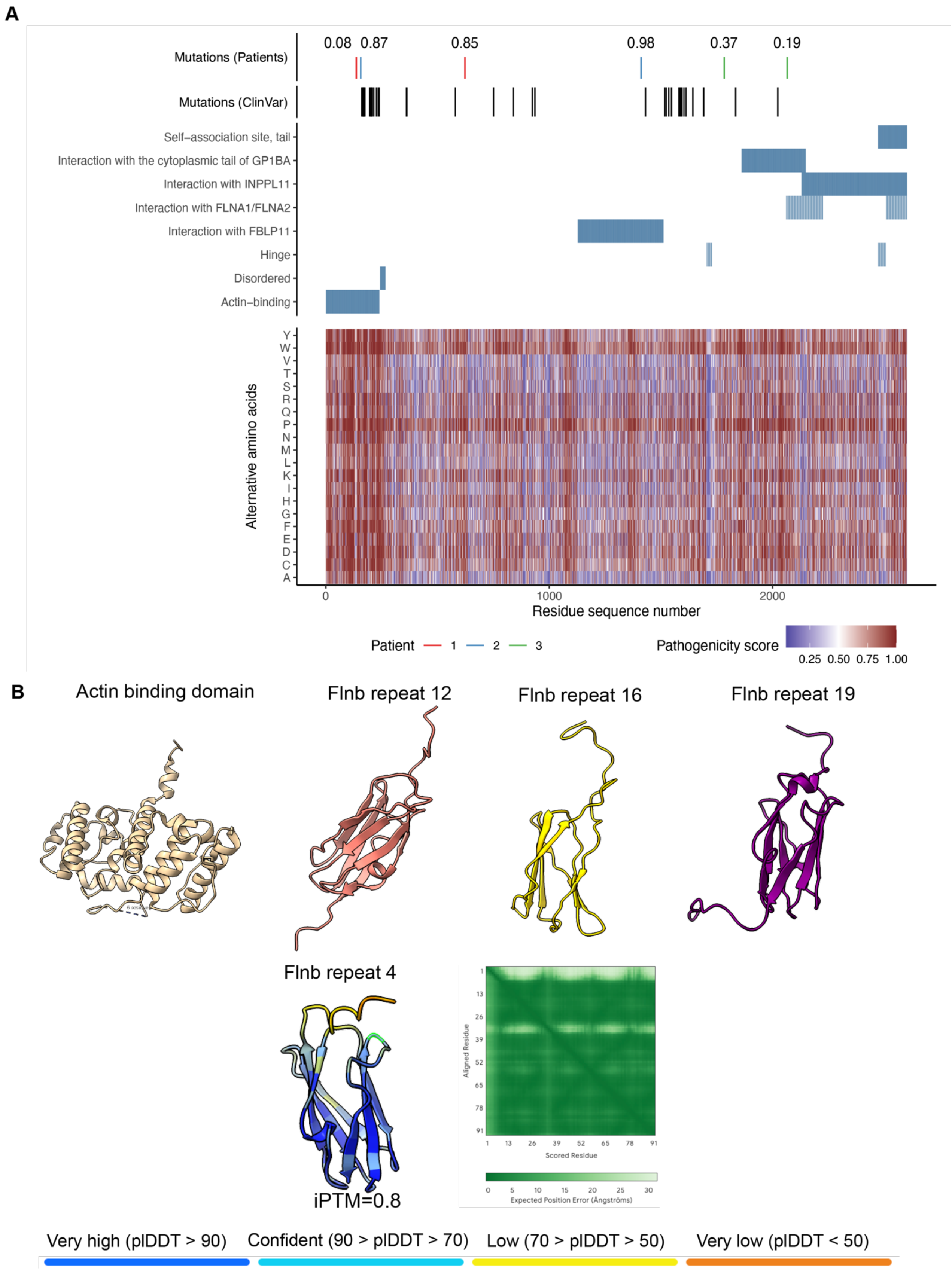


Figure S1

**Figure S1. AlphaMissense predictions for FLNB missense variants reveal pathogenic hotspots**

A. AlphaMissense was used to assess the predicted impact of recessive missense variants in three heterotaxy probands harboring FLNB mutations. At least one variant per proband was predicted as non-benign (pathogenicity score > 0.34). A comprehensive in silico analysis of all possible FLNB missense variants showed that over 80% of mutations in the actin-binding domain and disordered region were predicted to be pathogenic. AlphaMissense analysis of ClinVar-classified likely pathogenic and pathogenic FLNB missense variants associated with Larsen syndrome and other FLNB-related disorders revealed clustering within the actin-binding domain and the region between the Hinge and FBLP1-interacting domains. These patterns support a model in which specific FLNB domains are highly sensitive to missense perturbations.

B. The N-terminal Actin binding domain in the wild-type Flnb (RSCB PDB no. 2WA5), Filamin B repeat 12 (amino acids 1325-1422; RSCB PDB no. 2DIC)), FilaminB repeat 16 (amino acids 1726-1823; RSCB PDB no. 2EE9), Filamin B repeat 19 (amino acids 1999-2096; RSCB PDB no. 2DI8). There was no experimentally resolved structure available for the Filamin B repeat 4, so the corresponding region (from amino acids 544-636) was modelled using alphafold3.

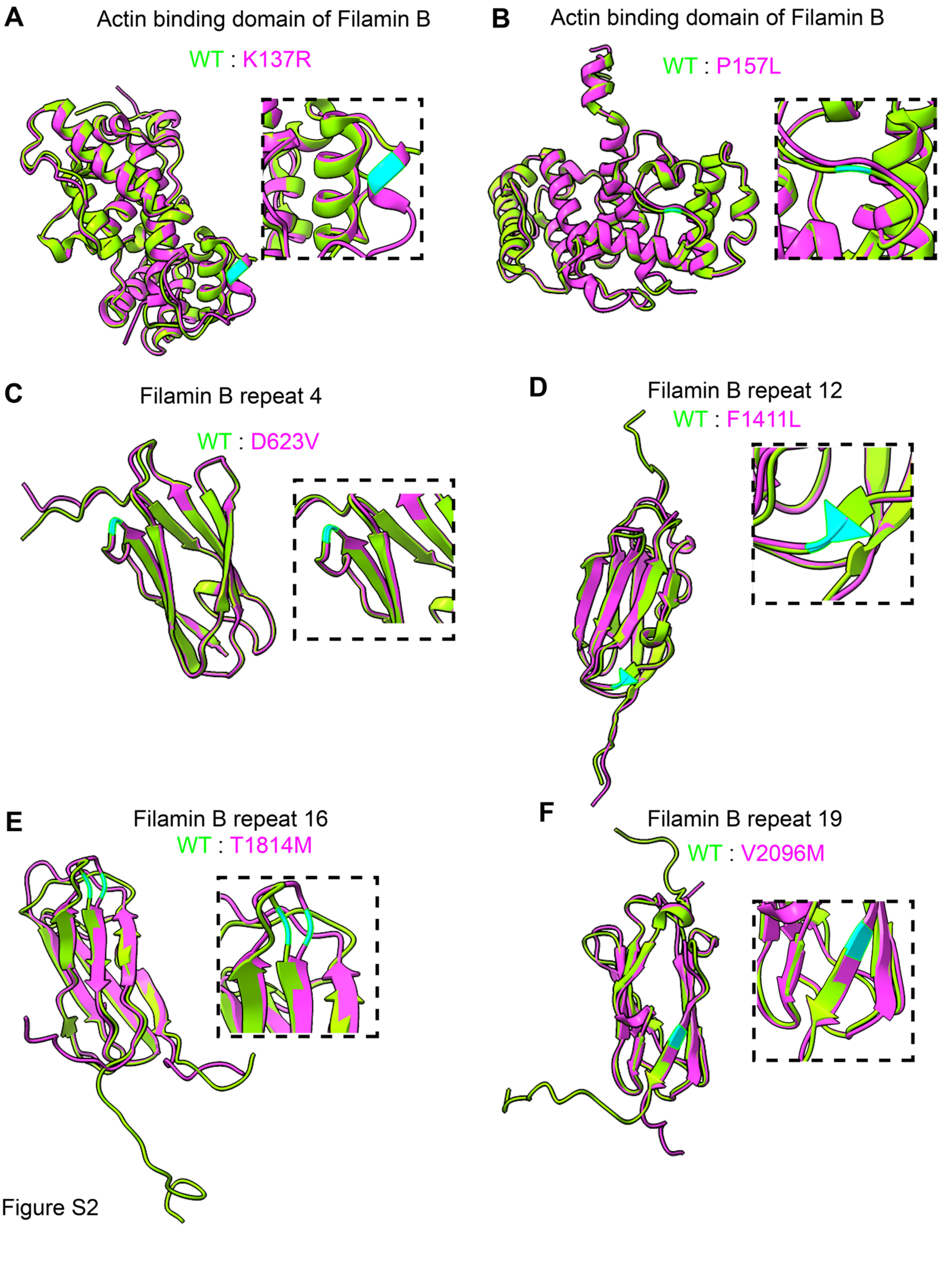
A
Actin binding domain of Filamin B
WT : K137R
B
Actin binding domain of Filamin B
WT : P157L
C
Filamin B repeat 4
WT : D623V
D
Filamin B repeat 12
WT : F1411L
E
Filamin B repeat 16
WT : T1814M
F
Filamin B repeat 19
WT : V2096M

Figure S2

**Figure S2. Comparison of the wild-type Filamin B structure and structures derived from patient mutations.**

A., B., the N-terminal actin-binding domain in the wild-type (RSCB PDB no. 2WA5; green) has two patient-derived mutations modeled using Alphafold 3 (K137R, P157L; magenta).

C. Comparison of Filamin B repeat 4 wild type (green) and the patient mutation D623V (magenta). Both structures were generated using AlphaFold3.

D. Comparison of Filamin B repeat 12 wild type (RSCB PDB no. 2DIC; green) and the patient mutation F1411V (or F1342V) (magenta). Both structures were generated using Alphafold 3.

E. The T1814M (T1783M isoform 2) patient mutation (magenta) is located in the Filamin B repeat 16 and is compared with the wild-type structure (RSCB PDB no. 2EE9; green).

F. The V2096M (V2065M isoform 2) patient mutation (magenta) is located in the Filamin B repeat 19 and is compared with the wild-type structure (RSCB PDB no. 2DI8; green). Mutation sites are labeled in cyan. The inset shows a zoomed-in view of the mutated site.

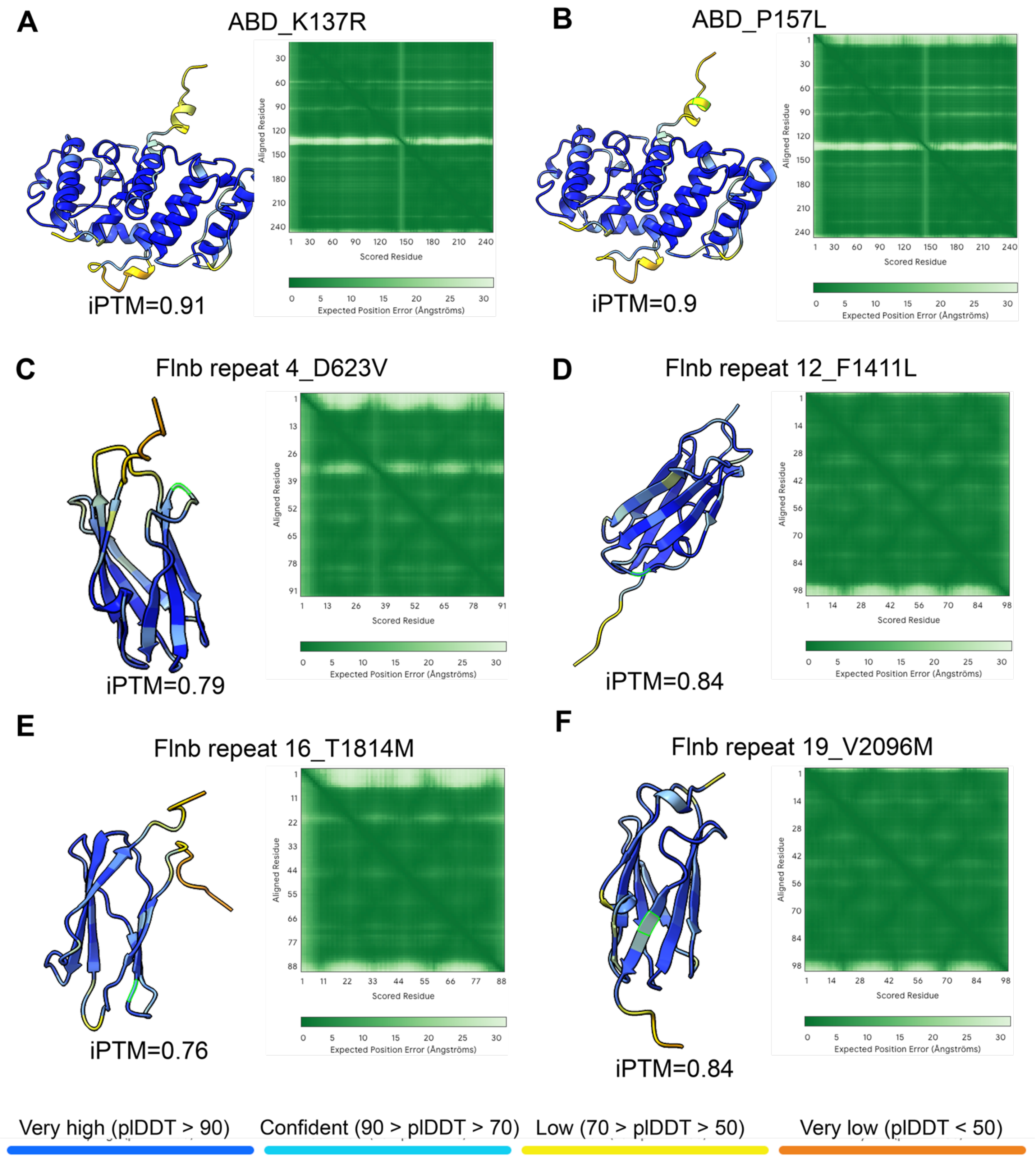
A
ABD_K137R
iPTM=0.91
B
ABD_P157L
iPTM=0.9
C
Flnb repeat 4_D623V
iPTM=0.79
D
Flnb repeat 12_F1411L
iPTM=0.84
E
Flnb repeat 16_T1814M
iPTM=0.76
F
Flnb repeat 19_V2096M
iPTM=0.84
Aligned Residue
Scored Residue
Expected Position Error (Ångströms)
Very high (pLDDT > 90)
Confident (90 > pLDDT > 70)
Low (70 > pLDDT > 50)
Very low (pLDDT < 50)

Figure S3

**Figure S3. Alphafold 3 modelled structures of patient derived FLNB mutations.**

The structures predicted are colored based on their pLDDT scores.

A-B. The N-terminal Actin binding domain 2 patient derived mutations (K137R, P157L). The N-terminal region from amino acids 1-242 of Filamin B was used for modelling.

C. Structure of patient mutation D623V in the Filamin B repeat 4 domain.

D. The F1411V (F1342V isoform2) patient mutation lies in the Filamin B repeat 12.

E. The T1814M (T1783M isoform 2) patient mutation lies in the Filamin B repeat 16.

F. The V2096M (V2065M isoform 2) patient mutation lies in the Filamin B repeat 19.

### SUPPLEMENTAL TABLE LEGEND

**Table S1.** Eighteen candidate genes were selected for CRISPR-mediated loss-of-function screening in *Xenopus* embryos based on predicted ciliary associations derived from analyses of eight databases.

| Gene | HPA | Cilia Miner | Cilia Carta | GeneCards | Syscilia | Primary Cilium | UCSC | Cildb | Total |
|---|---|---|---|---|---|---|---|---|---|
| FLNB | No | No | No | Yes | No | No | Yes | Yes | 3 |
| MPP4 | No | No | No | Yes | No | No | Yes | Yes | 3 |
| MYO15A | No | No | Yes | Yes | Yes | No | No | No | 3 |
| DYSF | No | No | No | Yes | No | No | No | Yes | 2 |
| EEA1 | No | No | No | Yes | No | No | No | Yes | 2 |
| ETS1 | Yes | No | No | No | No | No | No | Yes | 2 |
| FRYL | No | No | No | Yes | No | No | No | Yes | 2 |
| PPARA | No | No | No | Yes | No | No | No | Yes | 2 |
| ZC3H14 | No | No | No | Yes | No | No | No | Yes | 2 |
| ALS2 | No | No | No | No | No | No | No | Yes | 1 |
| FAM151B | No | No | No | No | No | No | No | Yes | 1 |
| PIWIL1 | No | No | No | No | No | No | No | Yes | 1 |
| DISP1 | No | No | No | No | No | No | No | No | 0 |
| GJA10 | No | No | No | No | No | No | No | No | 0 |
| LZTR1 | No | No | No | No | No | No | No | No | 0 |
| MYORG | No | No | No | No | No | No | No | No | 0 |
| UBR1 | No | No | No | No | No | No | No | No | 0 |
| ZNF704 | No | No | No | No | No | No | No | No | 0 |
